# Dynamic Inference by Model Reduction

**DOI:** 10.1101/2023.09.10.557043

**Authors:** Matteo Priorelli, Ivilin Peev Stoianov

## Abstract

How can agents infer the intentions of others by simply observing their behavior? And how can they generate fast and accurate actions such as grasping a moving object on the fly? Recent advances in Bayesian model reduction have led to innovative, biologically plausible approaches to actively infer the state of affairs of the world and perform planning with continuous signals. However, reducing the surrounding environment into a small set of simpler hypotheses remains a challenge in highly dynamic contexts. In this study, we propose an approach, based on active inference, that employs dynamic priors sampled from reduced versions of a generative model. Each dynamic prior corresponds to an alternative evolution of the world, which the agent can evaluate by accumulating continuous data. We test our approach on two everyday tasks: inferring a trajectory and grasping a moving object. Our findings reveal how agents can smoothly infer and enact dynamic intentions, and emphasize the key role of intentional gain or precision in motor learning.

## 1 Introduction

How can intelligent agents construct dynamic representations of the world that support high-level planning, control, and decision making? A recent brain-inspired framework, known as *active inference*, proposes that in order to survive, all organisms are constantly involved in minimizing a quantity called *free energy* [1, 2, 3, 4]. To remain within viable states, agents are supposed to maintain an internal generative model about how hidden states generate sensory observations and how such states evolve. The resulting predictions are compared with the observations to form prediction errors, which are minimized either by updating the agent’s beliefs (giving rise to perception) or by changing the world through action [5, 6].

Dealing with a continuous environment is possible by expressing the hidden states in terms of generalized coordinates consisting of multiple temporal orders (e.g., position, velocity, acceleration, and so on); this permits representing the environment dynamics with high fidelity [7, 8, 9], but only minimizes the free energy of present states and observations. In other words, simply responding reflexively to changes in sensory input precludes planning. This is sometimes referred to as “merely reflexive” active inference (e.g., as a thermostat or small insect). Decision-making is possible by augmenting a continuous model with a discretized representation of the environment, encoding possible future states. Planning then arises when future outcomes are treated as hidden states – an approach called *planning as inference* [10, 11, 12] – in which actions are selected to minimize the free energy expected in the future in consequence of that action [13, 14, 15]. However, selecting among a number of discrete courses of action (i.e., sequential policies) raises a problem: how does one infer which discrete trajectory is being pursued, given observations of continuous movement. In active inference, this is usually implemented using Bayesian model reduction to evaluate the evidence for a small number of discrete alternatives. Bayesian model reduction [16] is an efficient form of Bayesian model selection that has been used in several areas over the course of the years [17, 18, 19, 20, 21, 22, 23, 24]. This technique consists of reducing a complex distribution into a small set of simpler hypotheses or reduced models that have the same likelihood and differ only in their priors.

Hybrid models that integrate discrete and continuous components have been applied to several domains, ranging from epistemic foraging in reading, [25, 26], multi-step reaching movements in neurological disorders [27], saccades and visual sampling [28, 29], active listening [30], and interoceptive control [31]. However, the reduced models used for evaluating the full complex distribution are usually shaped a priori. This limits the agent to a static hypothesis space, making it difficult to adapt quickly in dynamic environments.

Here, we propose an alternative method that combines discrete hypotheses with potential trajectories. In summary, we address a fundamental problem in neuroscience and neurorobotics; namely, inferring the discrete intentions of another – or indeed oneself – based on patterns of continuous movement. This is relevant, not just in terms of action observation, but also in refining predictions of one’s own actions in the context of active inference. Technically, we solve this problem by combining Bayesian model reduction with a mixture of Gaussians over movement trajectories in generalized coordinates of motion. This allows the agent to flexibly construct and evaluate potential futures in real-time, enabling both efficient inference and rapid adaptation in constantly changing settings. This approach can be used in several dynamic contexts, and we demonstrate its capabilities in inferring which – of a number of moving targets – an agent is tracking, and grasping a moving object.

## 2 Results

### 2.1 Dynamic hybrid models

As described in Section 4.4, conventional hybrid models in active inference operate under the assumption that discrete hypotheses correspond to fixed priors over continuous latent states. In simple environments with little uncertainty this is often sufficient; however, it becomes limiting in real-world scenarios where intentions unfold dynamically over time, such as inferring a followed target or grasping a moving object.

The first problem can be addressed by considering the target positions as priors of reduced models, and comparing them with the posterior over hidden causes inferred by continuous data. However, the agent may fail to infer the causes of movement if the targets are constantly in motion, because the reduced models cannot be updated sufficiently quickly. A similar problem arises in the second example: in this case, a discrete model is needed to infer the correct sequence of actions needed to grasp the object, i.e., reach and then grasp. Again, the agent would fail the task if the object is moving too fast. Further, the object might also move during the grasping phase, so the agent has to carefully balance both actions depending on the object position at each continuous instant. The period of evidence accumulation actually poses a limit on how frequently the model can change the priors that must be realized or inferred.

In such cases, the agent’s set of hypotheses must evolve in accordance with the external world. This raises a key question: how can agents perform discrete inference and planning when the properties of the environment are constantly changing? Here, we address this issue by treating discrete outcomes directly as models of trajectories and not fixed set points. Such trajectories can be seen as hypothetical evolutions of the external world, which can be used either for inferring the actual evolution (e.g., related to the intentions of the self or other agents) or for imposing a particular evolution *to* the world (i.e., achieving dynamic goals by acting).

Formally, we want to construct a flexible discrete representation of the environment modeled by the following system of stochastic differential equations:

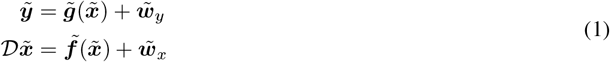

Here,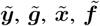 are respectively the continuous observations, likelihood function, continuous hidden states, and dynamics function, while 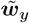 and 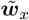 are noise terms sampled from Gaussian distributions, all encoded in generalized coordinates of motion – see Section 4.1 for more details.

We make use of the following hybrid generative model, displayed in Figure 1a:

**Figure 1.**
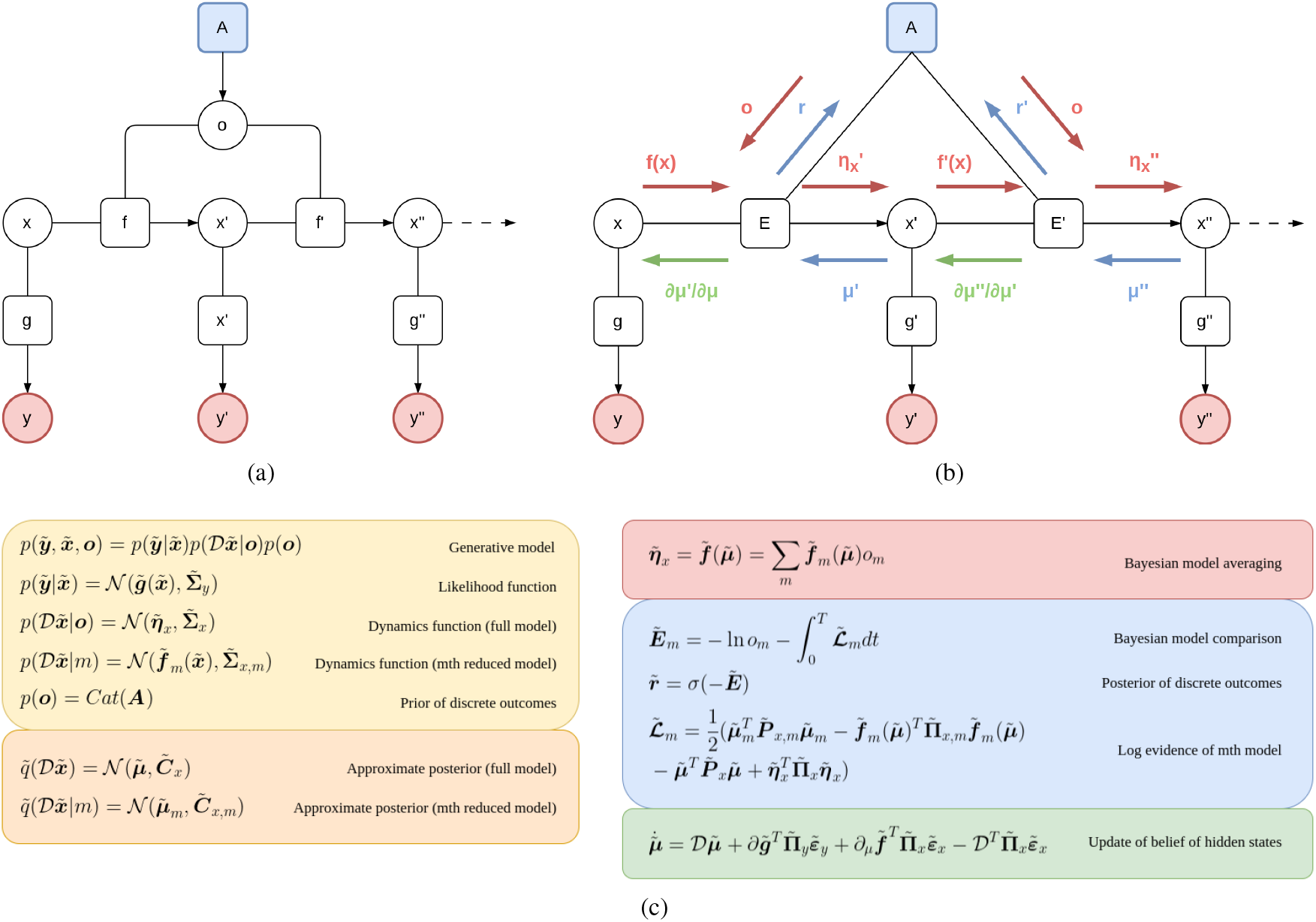
(a) Factor graph of a dynamic hybrid model. Discrete outcomes ***o*** encoding a dynamic set of hypotheses are generated by a prior ***A***. A potential evolution of the hidden states 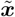 is generated by a dynamics function 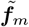, related to the *m*th dynamic hypothesis. In turn, the hidden states generate sensory predictions via the likelihood function 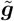, which are then compared with the corresponding sensations 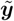. (b) Graph exemplifying the exchange of messages within a dynamic hybrid model. The full prior 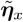 represents an average trajectory that depends on the potential trajectories and the discrete outcomes. In turn, the gradient 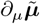 indicates how the average trajectory is mapped back into previous orders. Last, the posterior 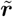 conveys the most likely agent’s hypothesis that could have generated the current trajectory, wherein each temporal order refines the estimation. This is done by computing the free energy of every reduced model and temporal order, i.e., ***E, E***^*′*^, and so on. (c) Generative model, approximate posteriors, Bayesian model averaging, Bayesian model comparison, and update rule of the beliefs.

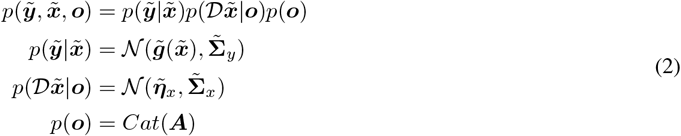

where ***A*** is a prior. In particular, the likelihood and dynamics are encoded with Gaussian probability distributions. Instead, discrete outcomes ***o*** are generated by a categorical distribution, wherein each element represents the probability that a specific continuous trajectory will be perceived, i.e., ***o*** = [***o***_1_, …, ***o***_*M*_], where *M* is the number of dynamic hypotheses. Further, we consider the distribution of the dynamics 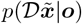 as a full model over the set of hypotheses, where 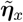 is the generalized prior over the trajectory. We can infer the most likely discrete outcome under the prior ***A*** if we assume that the agent maintains *M* reduced models:

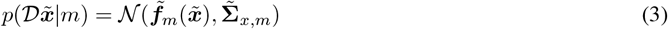

with the same likelihood as the full model but differing in their prior. Here, 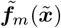 is the dynamics function of the *m*th reduced model. Following the variational approach described in Section 4.3, we define the corresponding approximate reduced and full models:

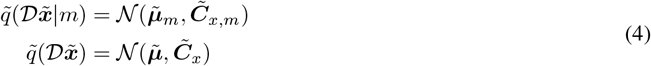

The implications of Equations 3 and 4 can be better understood from Figure 1b, which highlights three different kinds of message passing between discrete and continuous representations. Top-down messages are computed through a Bayesian model average of the prior trajectories 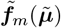 of each outcome model and the corresponding probabilities:

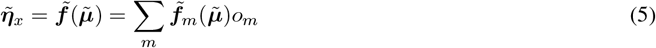

As a consequence, the agent does not maintain fixed priors over the environment, but dynamic priors that depend on potential trajectories and evolve at each continuous time step.

Concerning the inference, each reduced model is scored as in conventional hybrid models, but now the posterior over the current trajectories is compared with the trajectory generated by the dynamics functions of the hidden states:

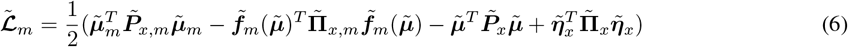

expressed in terms of posterior and prior precisions 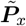 and 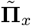. Here, the posterior mean and precision of each reduced model are:

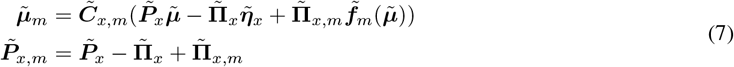

At this point, for each temporal order and model, a Bayesian model comparison is performed between the prior surprise and the log evidence accumulated over a time *T*. Finally, a softmax function is applied to get a proper probability:

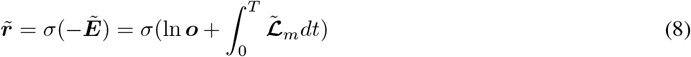

In summary, what Equation 5 means is that the agent has some dynamic hypotheses over the possible causes that could have generated its perceptions. It hence makes a guess by combining the related potential trajectories. The other side of the flow works similarly: the agent accumulates, for a certain amount of time, a score – i.e., the log evidence 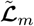 – of such dynamic hypotheses depending on continuous data, that is, it infers the most likely model that could have produced the perceived trajectory. The posterior 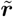 over discrete outcomes – expressed by Equation 8 – defines the agent’s final guess after observing the evidence, where each temporal order further refines the estimation. Since the potential trajectories are continuously generated, a model can be inferred that corresponds to a dynamic path for the whole period of evidence accumulation.

In relation to the backward pass – highlighted in green in Figure 1a – the prediction error over dynamics between the actual trajectory and the full prior, i.e., 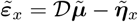, is backpropagated via the gradient of the dynamics functions with respect to the hidden states:

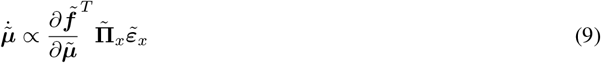

This gradient encodes how an average trajectory is mapped back to the current one. Specifically, for each temporal order the most likely continuous state that could have been generated from the previous order is inferred.

This architecture has a key implication related to the tasks mentioned above. If *o*_*m*_ is set to 1 and all the others to 0, the hidden states will be subject only to the *m*th trajectory. Due to the dualism between action and perception, if multiple outcome models are active, the hidden states will be pulled toward a combination of them, in the same way that a high-level policy generates the transitions for the discrete hidden states below. In other words, discrete outcomes not only represent the agent’s beliefs about the environment but also its intentions, now encoded with dynamic representations. As a result, a specific reaching movement can be found by generating and comparing dynamic hypotheses depending on the trajectories of the hand and every target, all inferred from continuous data. Additionally, grasping a moving object is possible by linking the prior to a discrete model, i.e., ***o***_*τ*_ = ***As***_*τ*_, where ***A*** is the likelihood matrix and ***s***_*τ*_ are the discrete hidden states at time *τ* – see Section 4.2. This process unfolds into the following steps: (i) the discrete model predicts an outcome corresponding to the reaching action; (ii) the discrete outcome steers the lower-level dynamics by generating a specific trajectory; (iii) this trajectory backpropagates in terms of dynamics prediction errors, eventually generating proprioceptive predictions needed for movement; (iv) after a period *T*, the discrete backward pass infers the most likely discrete outcome based on the accumulated evidence – i.e., it signals whether the hypothesis has been realized – and the discrete model eventually predicts the next (grasping) action. Because a discrete model can maintain a high-level intention for the whole period of evidence accumulation, a combined reaching-grasping action naturally emerges via inference – as we demonstrate in the simulations.

### 2.2 Two examples of dynamic inference

Ultimately, the choice of hybrid model depends on the task considered, with both types often necessary in realistic settings. Here, we briefly describe two examples where a dynamic hybrid model is fundamental. In the first one, the agent infers its intention based on the trajectories of its arm and two targets. The second example describes the control process of an agent aiming to reach and grasp a moving object.

#### 2.2.1 Trajectory inference

We first consider a situation in which an agent is reaching one of two moving targets and its goal is to infer which one it is following. The dynamic nature of the task can be tackled by defining potential trajectories for the objects of interest. Figure 2 shows a graphical representation of the hybrid generative model used by the agent. This is a basic inference task that does not involve high-level planning, thus we do not consider a complete discrete model. The agent controls an arm with a single Degree of Freedom (DoF). Its sensory modalities are: (i) a proprioceptive observation *y*_*p*_ for the arm’s joint angle; (ii) visual observations 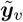 encoding the Cartesian positions and velocities of the hand and targets.

**Figure 2.**
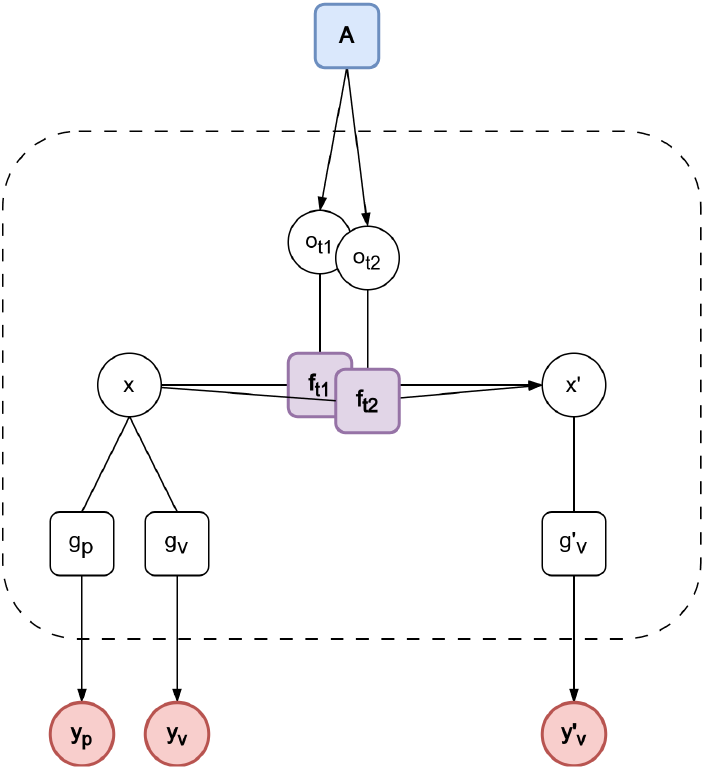
Factor graph for the trajectory inference task. Hidden states 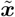 for the arm and the targets – encoded in joint angles – generate proprioceptive and visual predictions via likelihood functions *g*_*p*_ and 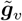, respectively. Through Bayesian model comparison, the most likely agent’s intention – encoded as discrete outcomes ***o*** – is dynamically inferred.

Since encoding Cartesian positions is not strictly necessary for this task, we express hidden states ***x*** in joint angles only, which thus generate and integrate both sensory modalities. The hidden states encode information about all the three elements of the environment:

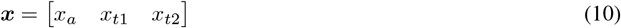

while the successive order ***x***^*′*^ expresses their angular velocities. Proprioceptive predictions – needed to infer the kinematic configuration of the agent’s body – are generated by a function *g*_*p*_ that simply extracts the arm’s joint angle from the belief 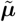 of the hidden states:

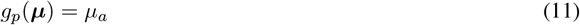

Instead, visual predictions are generated by a function 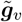 that computes the Cartesian positions and velocities of every component of the hidden states:

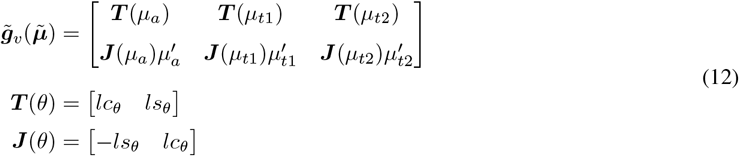

where ***T*** is the forward kinematics, ***J*** its Jacobian, *l* is the arm length, while *s*_*θ*_ and *c*_*θ*_ are the sine and cosine of the angle.

The prediction errors are then computed by comparing predictions with the corresponding observations:

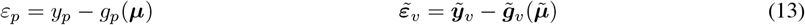

Discrete outcomes ***o*** – which we let here depend on a uniform prior – have only two elements, corresponding to the agent’s intentions to follow the first or the second target:

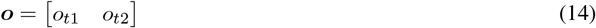

The belief dynamics can be quite complex and take into account many factors including multiple temporal orders, friction, gravity, and so on, but in the following we assume that it depends on a combination of these intentions and that the agent is velocity-controlled. We thus define two intentional future states in which the first component (corresponding to the arm’s joint angle) is equal to the inferred angle of the objects, i.e., ***i***_*t*1_(***µ***) = [*µ*_*t*1_, *µ*_*t*1_, *µ*_*t*2_] and ***i***_*t*2_(***µ***) = [*µ*_*t*2_, *µ*_*t*1_, *µ*_*t*2_]. Next, we define a dynamics function for each reaching intention:

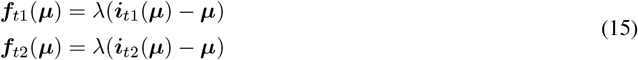

where *λ* is an attractor gain or precision. Such functions encode potential trajectories that the agent may observe, and correspond to attractors pointing toward a target – that is, the agent thinks that its arm will be pulled toward the inferred target angle. The dynamic prior 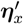 is found by weighting the potential trajectories with their corresponding probabilities that they will be perceived, i.e., *o*_*t*1_ and *o*_*t*2_. This is done through Equation 5, which is then used to compute a dynamics prediction error ***ε***_*x*_:

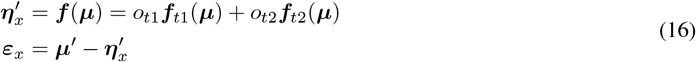

The belief of the hidden states is updated by minimizing the free energy:

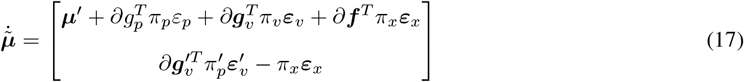

and using gradient descent. On the other hand, trajectory inference is done via Equations 6 and 8, where the posteriors over the reduced models are computed by Equation 7.

Figure 3 illustrates a simulation of the presented task. The arm and the two targets move at constant angular velocities. However, the red circle and the green square move slower and faster than the arm, respectively (see Figure 3a). As shown in Figure 3b, both discrete outcomes are initialized with equal probabilities, but the first outcome slowly increases as the arm approaches the first target. Then, it reaches a maximum, after which it starts to slowly decrease until the second outcome takes over. As explained before, the log evidence *L*_*m*_ of the *m*th reduced model scores how well the potential angular velocity of a target can explain the actual arm’s angular velocity (whose direction and magnitude are shown by the arrows in Figure 3a). The posterior ***r*** over discrete outcomes accumulate the evidence for the whole period in which the objects are moving, inferring the correct probability for each intention. Note that with an uninformative prior, Bayesian model comparison reduces to evidence accumulation. Overall, this simple task demonstrates that the model can discriminate between causes that generate continuous trajectories.

**Figure 3.**
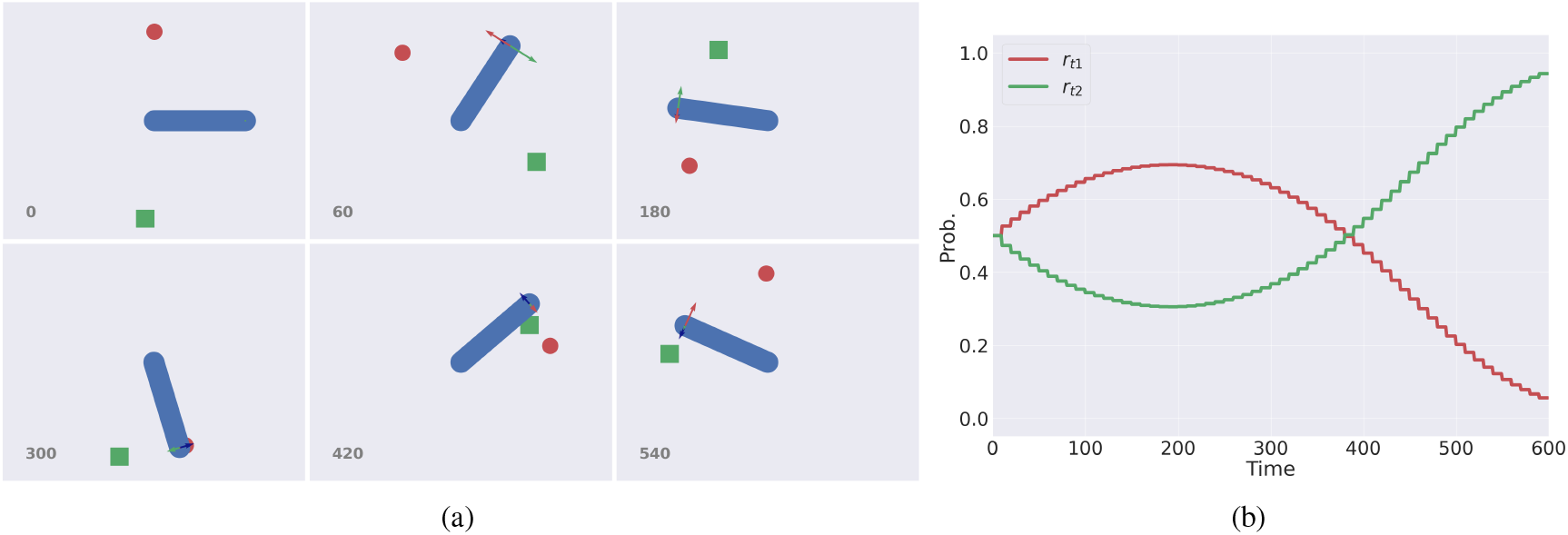
A simple inference task in which the agent (in blue) has to infer which one of two moving targets (a red ball and a green square) is following. See also [32] for a similar inference task in the Cartesian domain. (a) A sample sequence of time frames. The arm and the two targets move around the origin with constant but different velocities. The actual trajectory of the arm is represented with a blue arrow, while the potential trajectories of the two targets are displayed with red and green arrows. (b) Posterior over discrete outcomes, encoding the probability of reaching the red ball (*r*_*t*1_) and the green square (*r*_*t*2_), over time. The stepped behavior of the trajectories are due to the window *T* of continuous evidence accumulation of Equation 8, which was set to 10 continuous time steps.

#### 2.2.2 Object grasping

Grasping a moving object requires more complex computations than trajectory inference. In this case, the agent’s body is an 8-DoF arm where the last four joints correspond to the fingers. We use three observations: (i) proprioceptive observations ***y***_*p*_ = [*y*_*p*,1_, …, *y*_*p*,8_] encoding the joint angles of the arm; (ii) visual observations 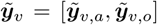 encoding the Cartesian positions and velocities of the hand and object; (iii) a discrete tactile outcome ***o***_*t*_ signaling whether the hand touches the object. We construct a discrete model with hidden states ***s*** encoding: (i) whether or not the hand is at the object position; (ii) whether the hand is open or closed; (iii) whether the hand has grasped the object or not. In total, these factors combine in 8 possible process states. The transitions encoded in the transition matrix ***B*** correspond to the three steps of the task, plus a *stay* action: (i) open the hand; (ii) reach the target; (iii) close the hand. The transition matrix is defined such that the object can be grasped only when both hand and object are in the same position.

We then define two dynamic hybrid models expressing intrinsic and extrinsic modalities (whose factor graphs are illustrated in Figure 4). Their generalized hidden states (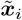 and 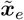) respectively encode information in intrinsic (e.g., joint angles and angular velocities) and extrinsic (e.g., Cartesian positions and velocities) coordinates. Moreover, both modalities comprise components corresponding to the arm and object, reflecting the decomposition of visual observations:

**Figure 4.**
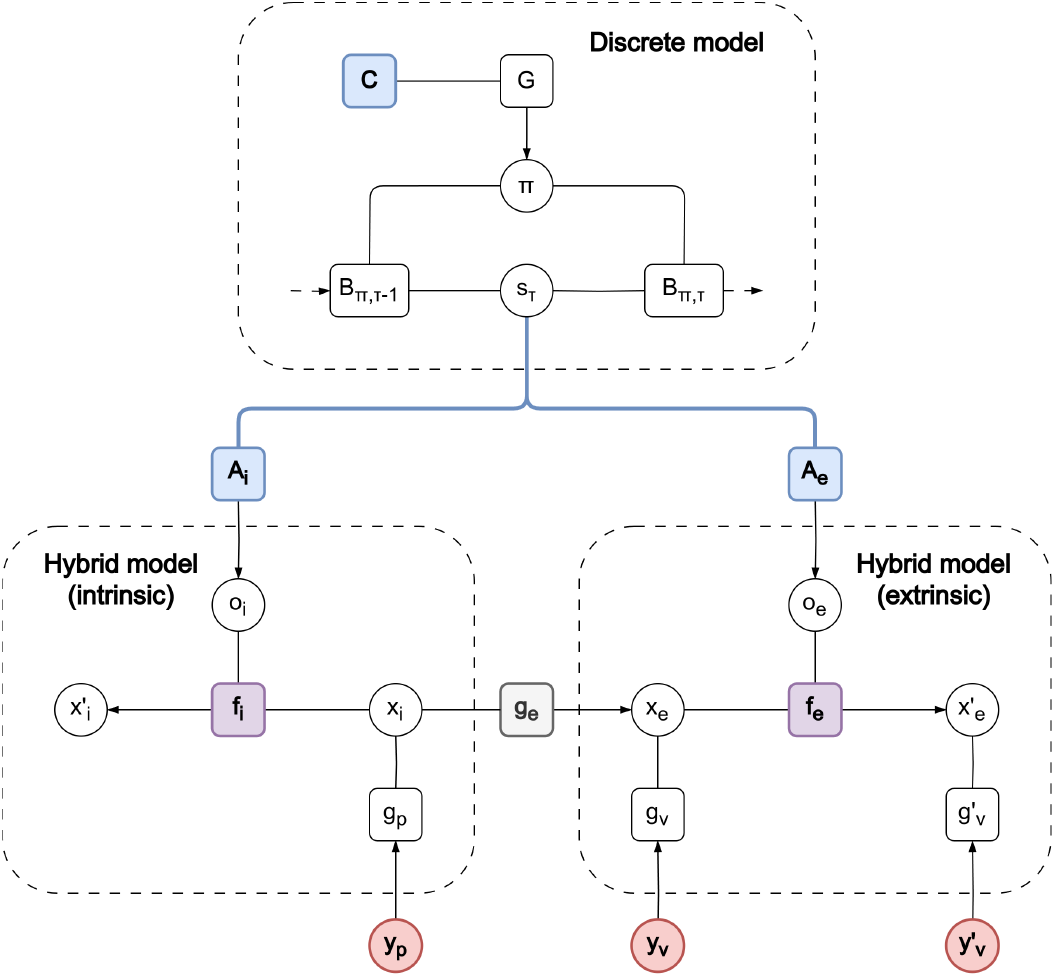
Factor graph for the object grasping task. Discrete hidden states ***s*** at time *τ* compute in parallel two predictions of discrete outcomes (***o***_*i*_ and ***o***_*e*_) for the intrinsic and extrinsic modalities via likelihood matrices ***A***_*i*_ and ***A***_*e*_. In turn, the discrete outcomes bias the continuous hidden states below (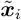 and 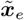), thus generating a particular trajectory. The two modalities are linked by a generative model ***g***_*e*_ that achieves kinematic inversion via inference, and they generate proprioceptive and visual predictions, respectively.

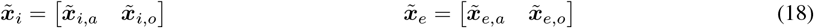

The 0th-order extrinsic hidden states ***x***_*e*,*a*_ and ***x***_*e*,*o*_ encode the positions of the hand and object, respectively. About the 0th-order intrinsic hidden states, while the interpretation of ***x***_*i*,*a*_ is intuitive (it just represents the actual body configuration related to the hand position), the other component can be seen as encoding a potential body configuration related to the object position [33]. In short, it may be considered as a state encoding a particular interaction with the object, depending on its affordances. The two modalities are linked via the 0th-order hidden states, by a likelihood function ***g***_*e*_ that performs forward kinematics:

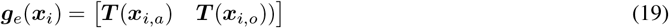

Since we are not interested in the fingers’ positions during the reaching task, these functions only compute the position of the middle of the two fingers. Maintaining a belief ***µ***_*e*_ in extrinsic coordinates greatly simplifies the computations because it permits expressing the dynamics without worrying about the inverse mapping to the joint angles, which is achieved by inverting the kinematic model via inference – for more details, see [34].

The discrete hidden states ***s*** simultaneously generate predictions for the hidden causes of intrinsic and extrinsic modalities through likelihood matrices ***A***_*i*_ and ***A***_*e*_. With this architecture, we can separately express grasping and reaching intentions in the two modalities:

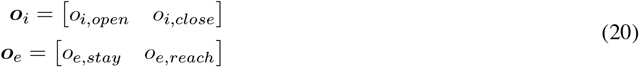

Nonetheless, more advanced movements – such as grasping an object with a specific wrist rotation – can be achieved by also expressing reaching intentions in the intrinsic modality [33]. Last, a matrix ***A***_*t*_ returns a prediction for the tactile observation ***o***_*t*_.

The discrete outcomes are linked to potential trajectories defining grasping and reaching intentions in the continuous domain. For the intrinsic modality:

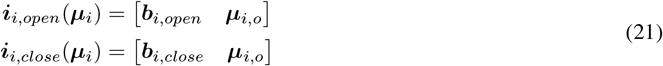

where ***b***_*i*,*open*_ and ***b***_*i*,*close*_ are the closed/open configurations. Correspondingly, for the extrinsic modality:

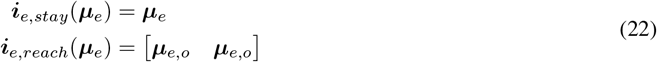

Here, ***i***_*e*,*stay*_ is an identity mapping and can be seen as an intention to maintain the current state of the world. Finally, we associate a dynamics function for each of these future states, i.e., ***f***_*i*,*open*_, ***f***_*i*,*close*_, ***f***_*e*,*stay*_, and ***f***_*e*,*reach*_.

The update of the hidden states follows the derivations of the previous example:

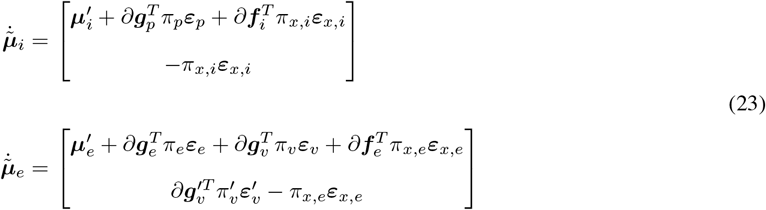

where ***g***_*p*_ and 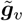 are simple identity mappings, while ***f***_*i*_ and ***f***_*e*_ are the functions computing a Bayesian model average of intrinsic and extrinsic trajectories. As concerns the posteriors over discrete outcomes at discrete time *τ*, they are updated by combining the agent’s surprise with continuous evidence:

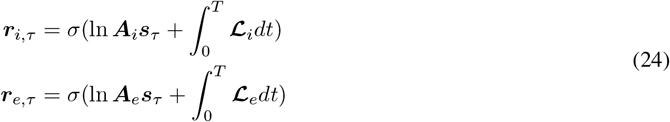

These messages are finally used by the discrete model, along with the tactile observation, to infer the most probable discrete hidden state at time *τ* and conditioned over policy *π*:

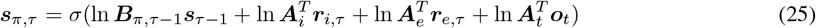

Note that the evidence accumulation is performed within the two dynamic hybrid models, so that the discrete model only needs to combine the two bottom-up messages, infer the most probable state, perform planning operations, and finally send predicted intentions back to the low levels, thus synchronizing their behavior.

Since the two hybrid models are now controlled by a discrete model, the agent can make a new plan if a grasping action fails because the object is moving too fast. This behavior can be seen in Figure 5b, which displays the trajectories of discrete hidden states ***s*** and intrinsic outcomes ***o***_*i*_. As soon as the hand approaches the object (at t=200), the agent makes a first grasping attempt, resulting in a gradual decrease of the hand-at-object state and a gradual increase of the closed-hand state. These states generate discrete outcomes in both modalities so that the *open* and *close* trajectories change accordingly. However, since the object is still too far from the hand, the agent rapidly turns back to tracking. At about t=400, the object is close enough to the hand, resulting in a much steeper change in the discrete hidden states and intrinsic outcomes. Eventually, the object is correctly grasped.

**Figure 5.**
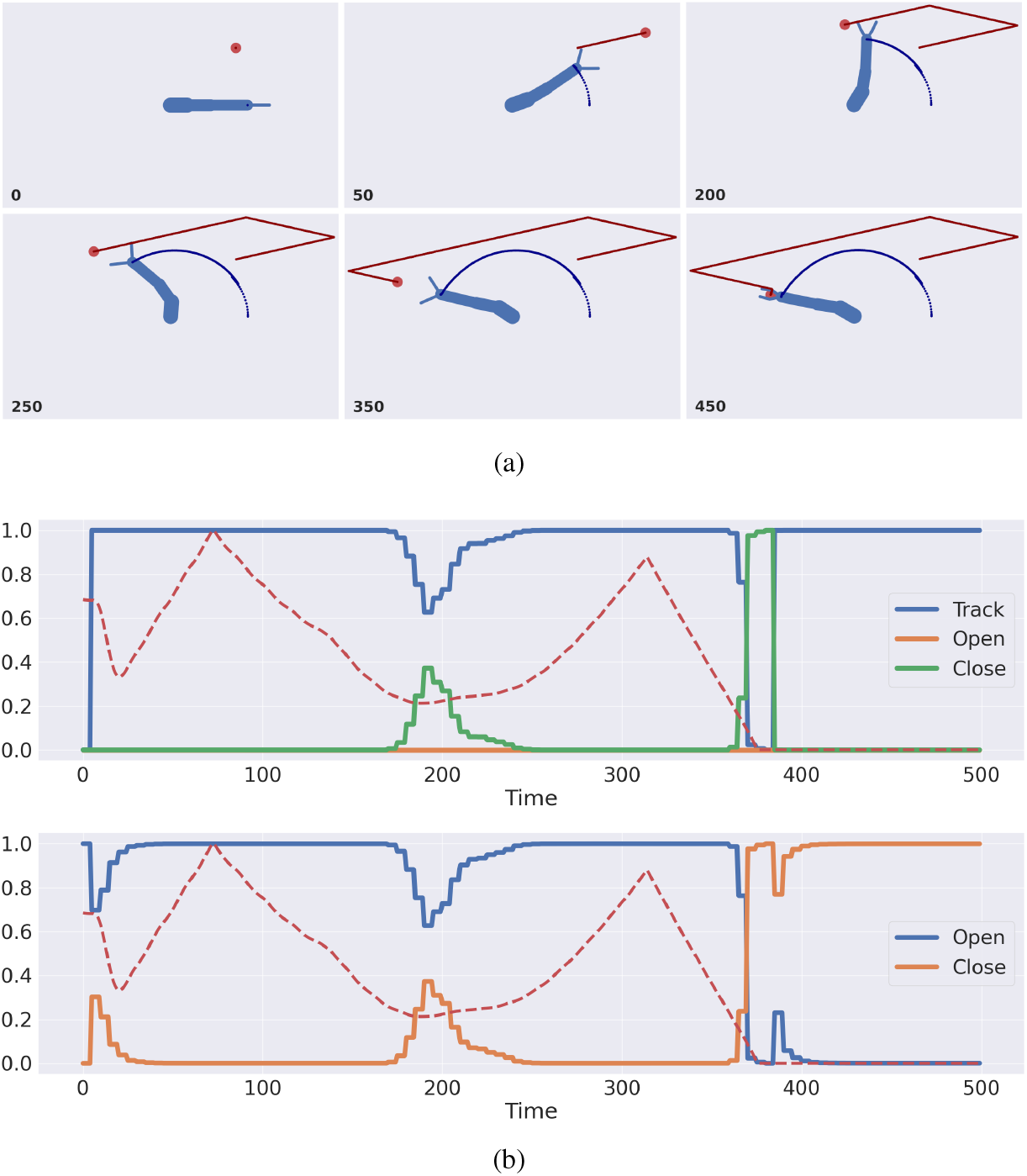
Object grasping with a 4-DoF arm and a 4-DoF hand with two fingers. (a) A sequence of time frames showing the arm (in blue) grasping the moving target (red ball). Trajectories of hand and object are shown, too. (b) Discrete hidden states ***s*** (top panel) and posterior over intrinsic discrete outcomes ***r*** (bottom panel), over time. Both panels also show the hand-object distance (dashed red line), normalized between 0 and 1.

## 3 Discussion

We presented a method that treats two fundamental problems – intention inference and motor control – under an active inference hybrid framework that relies on dynamic representations. The method leverages Bayesian model reduction [17], a powerful technique that, among its wide usages, allows shrinking continuous signals into a small set of hypotheses associated with discrete categories. In traditional active inference models, discrete outcomes are combined with static continuous priors to find the most likely hidden states based on the current evidence. This approach can be used, e.g., to reach fixed positions in sequence [1] or simulate pictographic reading [25]. However, using static assumptions limits applicability in real-world scenarios where goals and trajectories unfold dynamically. Our approach addresses this limitation by maintaining a set of dynamic hypotheses, that is, discrete outcomes related to potential trajectories. From the point of view of the continuous model, the discrete outcomes directly act as hidden causes, and a Bayesian model reduction is performed across multiple temporal orders of the hidden states.

This method permits drawing useful parallelisms on action understanding and policy evaluation. Specifically, top-down messages can be interpreted as the agent’s average intended trajectory (what the agent aims to realize), while bottom-up messages reflect the most likely intention that could have generated the observed movements. With this approach, evidence can be dynamically accumulated and associated with specific intentions. This allows the agent to constantly evaluate which potential evolution of the external world best explains its sensory observations, and act appropriately. This can be useful in every kind of context that presents dynamic elements, e.g., when interacting with moving objects.

An interesting fact regards the inference of self-generated movements. As also shown in [35], intention understanding in dynamic environments and embodied decision-making are two emergent properties of the proposed architecture. Although it may seem counterintuitive, inferring a decision from one’s own perceptions and movements is actually a key aspect of biological behavior that helps to reinforce and stabilize the decision taken [36, 37], ensuring that valuable opportunities are not missed in ecological settings. This mechanism has received little attention in artificial intelligence and robotics, but it could support advances in the next generation of intelligent agents. Finally, the presented model can be scaled up to control complex kinematic structures, synchronizing the behavior of multiple limbs and performing complex operations, e.g., for tool use [34, 38].

The interaction between inference and action becomes even clearer when we compare the precisions 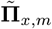 of the reduced dynamics functions of Equation 6 and the sensory precisions 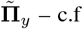., attentional gain or selection. In predictive coding, sensory precisions both encode the agent’s confidence about a sensory modality (e.g., when visual input is clear and unambiguous, visual precision is high), and attention (e.g., in a noisy environment, auditory precision may be low, so the agent shifts reliance to vision) [39]. In our case, a high precision assigned to a potential trajectory means it is a good option for minimizing the free energy or, more clearly, for inferring the current agent’s trajectory or realizing its goal. Conversely, a low precision implies that the trajectory is not useful for the task – e.g., a grasping action when the object is out of reach – or that it cannot be helpful to understand the situation – e.g., ignoring a target unrelated to the motion of another agent. In fact, attention is closely related to intention, in the sense that both are selected on the basis of relative precision or confidence. Attentional selection relates to state estimation, while intentional selection relates to action and execution. When considering learning of such precisions, two concurrent processes arise: a *fast* process (described here) that selects trajectories based on current goals and evidence; and a *slow* learning process that updates how useful each trajectory is in different contexts – in the same way that unreliable sensors get lower precision over time [40].

Overall, the model that we presented might be helpful to understand how dynamic hypotheses are encoded in a biologically plausible way. But it might also explain how intentions are assigned to specific tasks and how they emerge from high-level computations. As shown in Figure 5b, although we used elementary reaching and grasping actions, a smooth approach transition emerges, due to the Bayesian model reduction between potential trajectories. This novel behavior could be mapped into the continuous dynamics and called in a single action by the discrete model, producing a flexible and fluid behavior – as simulated in [33].

Future work may extend this approach in several directions. One limitation of this study is that we kept the precisions of the dynamics functions fixed. Learning the dynamics functions of the reduced models, along with their precisions, would be a promising direction of research to simulate task specialization and construction of richer behaviors from primitive movements. To address this issue, an alternative approach was implemented in [41], which used multiple continuous generative models followed by a “switcher”. This approach has some analogies with the model presented here in that the agent (a student bird) could maintain models of how teachers generate a possible evolution of sensory signals, and through the switcher it could perform online model selection to infer the most likely causes. Another study proposed a hybrid active inference model based on recurrent switching linear dynamical systems (rSLDS) [42]. This allowed the agent to learn discrete representations of continuous dynamics via piecewise linear decomposition, useful to solve complex tasks such as the Mountain Car problem.

Another limitation is the use of just two temporal orders, whereas trajectory inference with increasing temporal orders might be helpful to develop more precise robots or, for neuroscience, to decode intentions from gaze or other highly dynamic cues. Finally, the model could be used for multi-agent interaction, where an agent has to observe and imitate the intentions of other agents, either in collaborative or competitive settings.

## 4 Methods

### 4.1 Active inference in continuous time

Active inference is a brain-inspired theory that proposes a unifying paradigm to understand the behavior of all living organisms. It is based upon the *free energy principle* which states that, in order to survive, every creature must actively minimize the *free energy* [1, 2, 3, 4], which is the opposite of what is known in the machine learning community as *evidence lower bound* (or ELBO). The theory of active inference makes the same assumptions of predictive coding, i.e., that organisms perceive their environment through an internal generative model, inferring how external causes (of the real generative process) lead to the observed sensory signals [4, 2]. Regarding the continuous domain, this procedure is broken down into probability distributions comprising hidden causes 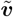, hidden states 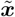, and sensory signals 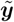:

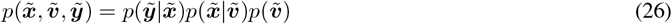

These distributions are approximated through Gaussian functions:

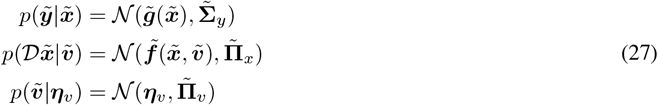

which respectively correspond to: (i) the likelihood function 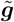, encoding how hidden states generate sensory signals; (ii) the dynamics function 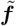, specifying the evolution of the hidden states; (iii) the prior ***η***_*v*_ over the hidden causes, whose role is to define targets for goal-directed behavior and to link hierarchical levels. Further, 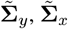, and 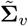 are the variances of sensory signals, hidden states, and hidden causes. Note that the symbol ~ denotes a variable in generalized coordinates of motion, encoding instantaneous trajectories up to a certain order, i.e. 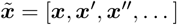, and that *𝒟* is a shift differential operator such that 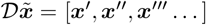. These functions are related to the following nonlinear stochastic equations that describe the evolution of the environment:

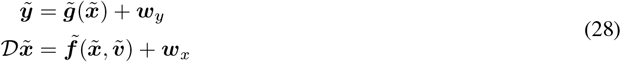

where 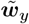 and 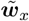 are noise terms sampled from Gaussian distributions.

In active inference, beliefs are updated not only by inferring the causes of sensory signals, but also by defining priors about their evolution over time. The dynamics of a hidden state does not necessarily need to be the same as the true generative process: indeed, it is this discrepancy that produces an action and makes the agent fulfill prior expectations. Belief dynamics can be seen as a combination of two components: (i) a model of how hidden states normally evolve based on past experience – e.g., expecting a moving ball to follow a straight path; (ii) a distorted vision of this dynamics that depends on the agent’s desires – e.g., thinking that my hand will be pulled toward the ball will drive the movements of my hand toward my expectations, even if that force does not exist in reality.

The agent’s goal is to compute the posterior 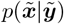, whose direct computation is however unfeasible, as it involves the calculation of the inaccessible marginal 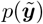. The variational approach involves approximating the posteriors with manageable distributions, such as the following Gaussians:

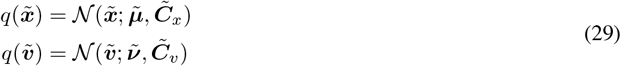

where the parameters 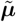 and 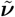 are the beliefs of hidden states and hidden causes, respectively. Perceptual inference then becomes a minimization process, aiming to reduce the disparity between the approximated and actual posteriors. This can be framed in terms of a KL divergence – equivalent to the expectation, over the approximated posterior, of the discrepancy between the two log probability distributions:

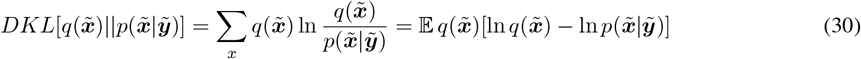

As the denominator 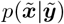 still depends on the marginal 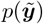, the KL divergence is expressed in terms of the log-evidence and the *Variational Free Energy* (VFE), with the latter being minimized instead. Given that the KL divergence is non-negative (due to Jensen’s inequality), the VFE serves as an upper bound on surprise. Consequently, minimizing the VFE enhances both the evidence and model fit:

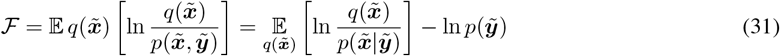

With the given approximations, the minimization of the free energy involves an iterative update of parameters through the following expressions:

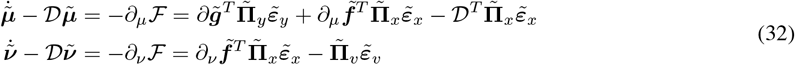

expressed in terms of precisions (or inverse variances) 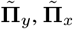, and 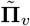. Here, 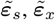, and 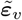 respectively denote the prediction errors of sensory signals, hidden states dynamics, and prior over hidden causes:

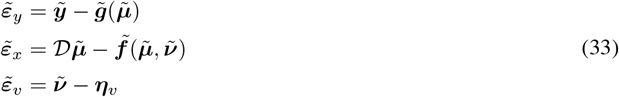

Importantly, the VFE can also be employed to compute motor control signals:

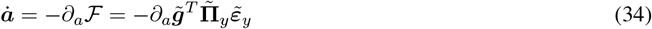

This process selects the outcomes that correspond to the representation of the world currently maintained by the agent; in short, if during perceptual inference the agent’s beliefs are updated so as to reflect the sensory signals, during *active inference* the agent acts with the goal of sampling those sensory signals that conform to its beliefs, as in a self-fulfilling prophecy. This sequence further minimizes the VFE, enabling the implementation of goal-directed behavior and maintaining the agent within predictable regimes of state space that are characteristic of the agent in question (e.g., avoiding collisions and reaching targets). [4].

### 4.2 Active inference in discrete time

While active inference in continuous time can address real-world challenges by tracking the instantaneous trajectories generated by the continuous environment, its applicability is constrained as it struggles to accommodate broader classes of actions and decision-making processes. The VFE minimization can solely update the approximate posterior based on current or past observations, failing to incorporate possible future states and outcomes. To grant this ability to the agent, a quantity called the *Expected Free Energy* (EFE) is introduced [13, 14]. We define an active inference discrete model as a Partially Observable Markov decision processes (POMDPs):

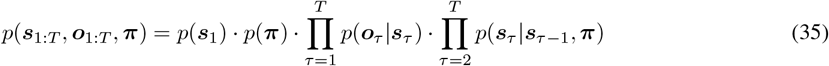

Here, ***s*** and ***o*** denote discrete states and outcomes. Notably, these policies are not mere stimulus-response mappings like those in Reinforcement Learning schemes; rather, they encompass sequences of actions. Given that action planning necessitates the selection of policies leading to desired priors, future outcomes not yet observed must also be taken into account. Hence, the EFE is formulated by conditioning over these unobserved outcomes, treating them as hidden states:

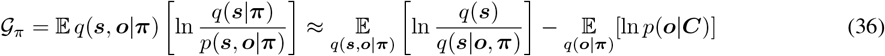

Here, ***s*** and ***o*** denote discrete states and outcomes, ***π*** the policies, *τ* a discrete time step. Notably, the policies are not mere stimulus-response mappings as in Reinforcement Learning schemes, but rather sequences of actions. Each of these elements can be expressed using categorical distributions:

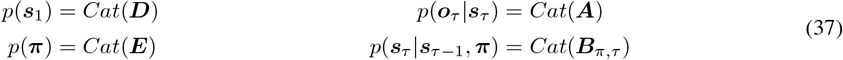

where ***D*** captures beliefs about the initial state, ***E*** encodes the prior over policies, ***A*** represents the likelihood matrix, and ***B***_*π*,*τ*_ is the transition matrix. As before, exact computation of the posterior *p*(***s***_1:*T*_, ***π***|***o***_1:*T*_) is unfeasible given the intractability of the model evidence *p*(***o***_1:*T*_). Therefore, we resort to an approximate posterior distribution *q*(***s***_1:*T*_, ***π***), and minimize the Kullback-Leibler (KL) divergence between the approximate and real posteriors:

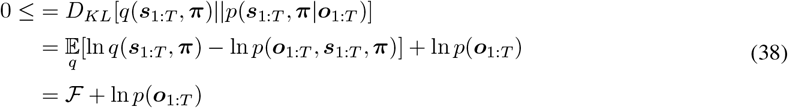

Assuming a mean-field approximation, the approximate posterior factorizes into independent distributions:

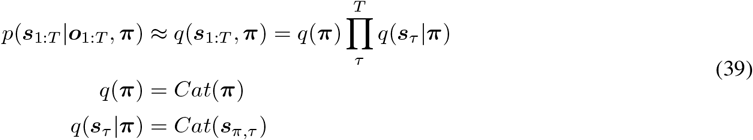

Variational message passing can then be employed to infer each posterior *q*(***s***_*τ*_ |***π***) and combine them into a global posterior *q*(***s***_1:*T*_ |***π***). To update the posterior about hidden states, the method combines messages from past and future states, as well as outcomes. This entails expressing each term in relation to its sufficient statistics and subsequently applying a softmax function to get a valid probability distribution:

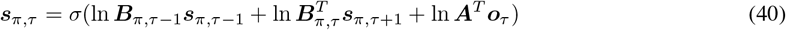

Likewise, to update *q*(***π***), the approach involves combining messages from the policy’s prior (expressed by matrix ***E***) and from future outcomes conditioned on policies – given that action planning requires the selection of policies that lead to desired priors, future outcomes not yet observed must also be taken into account. Hence, the EFE is formulated by conditioning over these unobserved outcomes, treating them as hidden states:

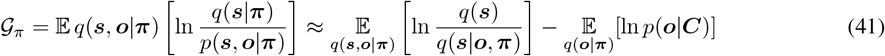

Here, the probability distribution *p*(***o***|***C***) represents preferred outcomes. The last two terms are referred to as *epistemic* (uncertainty-reducing) and *pragmatic* (goal-seeking) terms, respectively. Under appropriate assumptions, the EFE conditioned at a specific time step *τ* becomes:

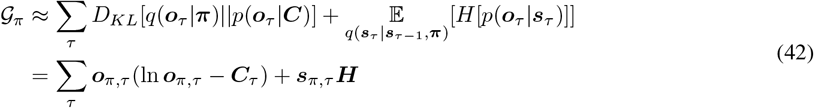

where:

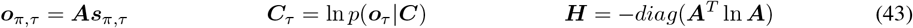

Ultimately, the most probable action under all policies is selected:

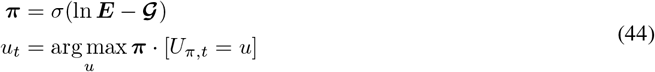

whose result in the same *self-evidencing* mechanism observed in the continuous framework.

### 4.3 Bayesian model reduction

Consider a generative model *p*(***θ, y***) with parameters ***θ*** and data ***y***:

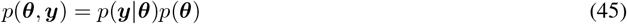

Consider then an additional distribution *p*(***y, θ***|*m*): this is a reduced version of the first model if the likelihood of some data is the same under both models and the only difference rests upon the specification of the priors *p*(***θ***|*m*). We can express the posterior of the reduced model in terms of the posterior of the full model and the ratios of the priors and the evidence:

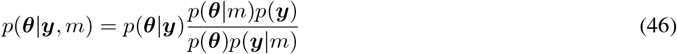

The procedure for computing the reduced posterior is the following: first, we integrate over the parameters to obtain the evidence ratio of the two models:

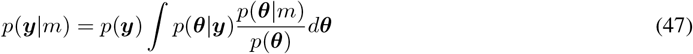

Then, we define an approximate posterior *q*(***θ***) and we compute the reduced Variational Free Energy (VFE) in terms of the full model:

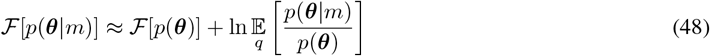

In other words, this quantity acts as a hint to how well the reduced representation explains the full model. Similarly, the approximate posterior of the reduced model can be written in terms of the posterior of the full model:

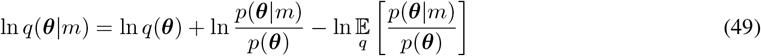

The Laplace approximation [43] leads to a simple form of the approximate posterior and the reduced free energy. Assuming the following Gaussian distributions:

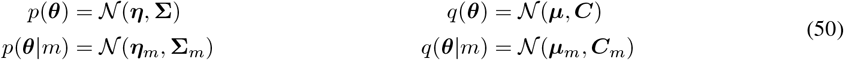

The reduced free energy turns to:

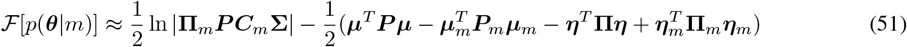

expressed in terms of precisions of priors ***P*** and posteriors **Π**. The reduced posterior mean and precision are computed through Equation 49:

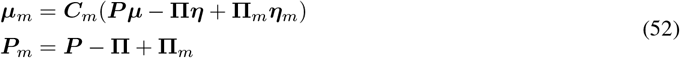

For a more detailed treatment on Bayesian model reduction, see [16, 17].

### 4.4 Hybrid models in active inference

Figure 6a shows the factor graph of a conventional hybrid model, whose generative model is assumed to be the following:

**Figure 6.**
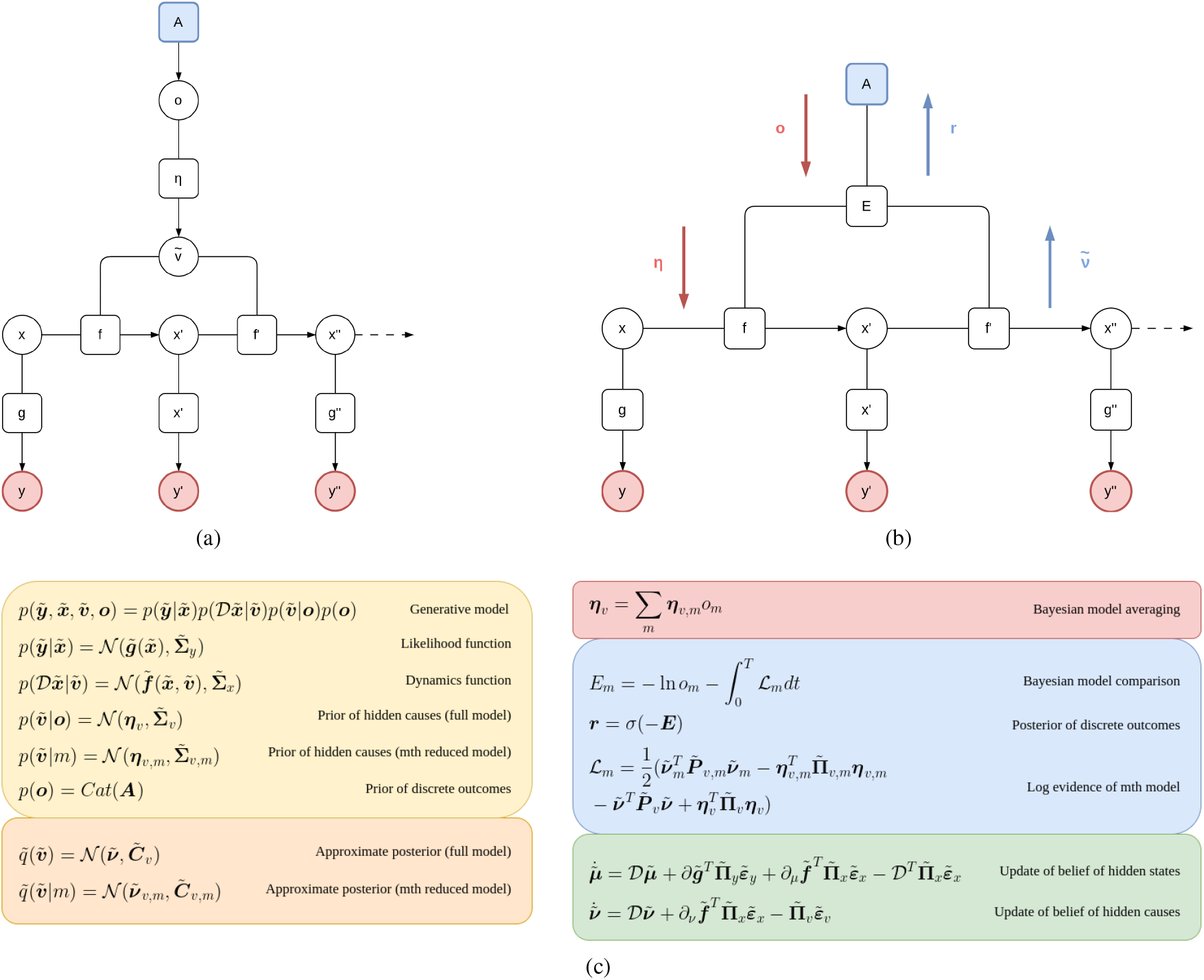
(a) Factor graph of a conventional hybrid model. Discrete outcomes ***o*** are generated by a prior ***A***, which we can assume coming from a discrete model with hidden states ***s*** – as in Section 4.2. In turn, the discrete outcomes act as a prior over the hidden causes 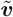 of the continuous model. (b) A graph highlighting the top-down (red) and bottom-up (blue) messages of Bayesian model reduction. The prior ***η*** is computed by Bayesian model averaging, which combines continuous priors ***η***_*v*,*m*_ of specific outcome models (not displayed here) with the respective probabilities *o*_*m*_. Bottom-up messages ***r*** entail a Bayesian model comparison, which compares the prior expectation of each model encoded in the surprise ln *o*_*m*_ with the respective log evidence ℒ_*m*_ accumulated over time. (c) Generative model, approximate posteriors, Bayesian model averaging, Bayesian model comparison, and update rule of the beliefs.

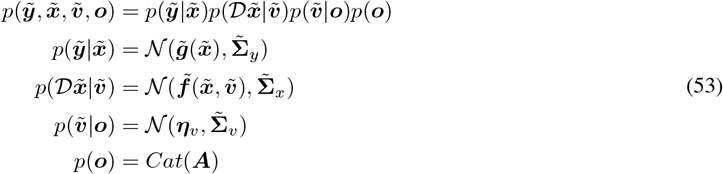

where 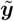 are the continuous observations, 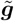 is the likelihood function, 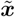 are the continuous hidden states, 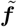 is the dynamics function, 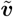 are the hidden causes, ***o*** are the discrete outcomes with prior ***A***. For simplicity, we do not consider a complete discrete model as in Section 4.2, but assume that discrete outcomes can be generated by discrete hidden states at time *τ* through the likelihood matrix, i.e., ***o***_*τ*_ = ***As***_*τ*_. This generative model is supposed to learn a representation of the environment modeled as in Section 4.1.

The probability distribution 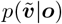 tells us that the hidden causes are generated by discrete outcomes ***o*** through a Gaussian distribution with mean ***η***_*v*_; this is considered a full model over a complex set of hypotheses. We can infer the most likely hidden cause under the current discrete outcomes by assuming that the agent maintains *M* reduced models:

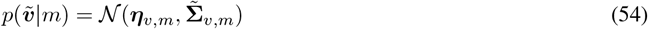

with the same likelihood of the full model but differing in their prior. This is a Gaussian mixture model, where the mixtures pertain to the generalized causes of a trajectory. In other words, we treat each outcome *o*_*m*_ as a particular model of dynamics, with prior ***η***_*v*,*m*_ over the hidden causes. As highlighted in Figure 6b, the exchange of top-down and bottom-up messages between the discrete and continuous representations follows the derivations of Bayesian model reduction in Section 4.3. After having defined appropriate approximate posteriors, the continuous expectations in Equation 54 are computed through a Bayesian model average between the continuous priors of each outcome model, weighted by the respective prior probabilities:

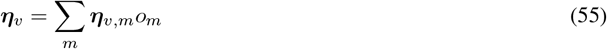

Similarly, the probability distribution ***r***, over models, is found by first computing a Bayesian model comparison between the prior surprise and the log evidence accumulated over a continuous time *T* :

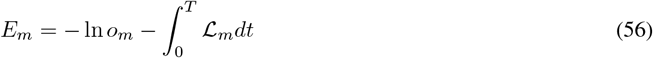

where *E*_*m*_ and *ℒ*_*m*_ are respectively the free energy and log evidence of the *m*th outcome model; then passing the free energy through a softmax function *σ* to get a proper probability, i.e., ***r*** = *σ*(*−****E***). The log evidence approximates the reduced free energy of Equation 51, where the posterior is over hidden causes 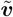:

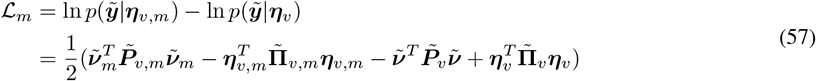

As a practical example, the reduced priors ***η***_*v*,*m*_ could encode hypothetical positions of the agent’s hand. Such reduced priors could be associated with different discrete hidden states and outcomes, encoding the probability that the defined positions will be observed (or realized). Via Equation 55, a discrete model can transform a high-level plan into a composite trajectory, e.g., for multi-step reaching [27]. Conversely, Equation 56 evaluates the probability of each reduced prior compared to the current posterior about the hand position. This message signals whether the agent has reached one of the positions, enabling the discrete model to compute the next reaching movement. See [25, 28] for more details on hybrid models.

## Acknowledgments

This research received funding from the European Union’s Horizon H2020-EIC-FETPROACT-2019 Programme for Research and Innovation under Grant Agreement 951910 to I.S. and the Italian Ministry for Research MIUR under Grant Agreement PRIN 2017KZNZLN to I.S. The funders had no role in study design, data collection and analysis, decision to publish, or preparation of the manuscript.

## Code availability

Code and data are deposited in GitHub: https://github.com/priorelli/dynamic-inference

